# Unraveling a fine balance between ferroptosis, lipid metabolism, and hormonal protection in Leydig cell steroidogenesis

**DOI:** 10.64898/2026.07.03.736405

**Authors:** Yanina Benzo, Melina A Dattilo, María A Raggio, Paula F Lopez, Denisse Ferrer Viñals, María S Theas, Cecilia Poderoso, Paula M Maloberti

## Abstract

Leydig cells (LCs) are essential for male reproductive function due to their role in testosterone synthesis, a process critically dependent on mitochondrial cholesterol transport mediated by the Steroidogenic Acute Regulatory protein (StAR). Despite their importance, LCs are highly sensitive to metabolic and exogenous stressors. Ferroptosis, an iron-dependent form of regulated cell death driven by lipid peroxidation, has emerged as a key link between cellular metabolism and cell fate; however, its role in LCs and steroidogenesis remains poorly understood. In this study, we investigated the induction of ferroptosis in LCs and its impact on their steroidogenic capacity. We evaluated cellular responses to canonical ferroptosis inducers (Erastin and RSL3) alongside the transcriptional regulation of key genes. Our results demonstrate that LCs are vulnerable to ferroptotic stress, which significantly downregulates *Star* expression. Notably, we uncovered a novel endocrine-metabolic crosstalk: hormonal stimulation via hCG effectively rescues LCs from Erastin-induced toxicity and fully sustains maximal steroidogenesis. However, this hormone-driven cytoprotection fails against direct GPX4 inhibition by RSL3, indicating an absolute reliance on functional GPX4. These mechanistic findings highlight the paradoxical dual role of ACSL4 in Leydig cell biology and are further supported by single-cell transcriptomic analysis of human testis, which reveals an imbalance in the ACSL4/GPX4 axis within the Leydig-cell compartment of infertile patients. Together, our data position ferroptosis as a critical disruptor of male endocrine function and reveal a hormone-mediated metabolic adaptation that could inform novel therapeutic strategies against oxidative stress in the testis.

**Highlights:**

– Leydig cells exhibit a strong vulnerability to ferroptotic cell death.
– Ferroptosis disrupts StAR expression and halts Leydig cell steroidogenesis.
– hCG signaling promotes metabolic adaptation against Erastin-induced ferroptosis.

## 1. Introduction

Leydig cells (LCs) are the primary steroidogenic cells in the testis, responsible for maintaining male fertility and systemic hormonal balance. Located in the testicular interstitial space, LCs are tasked with the production of testosterone (T), which is synthesized and immediately secreted upon stimulation by the luteinizing hormone (LH) from the anterior pituitary (Zirkin & Papadopoulos 2018). The first limiting-step of steroid production in every steroidogenic tissue is cholesterol transport into mitochondria, mediated by the Steroidogenic Acute Regulatory protein (StAR) (Privalle *et al*. 1983; Poderoso *et al*. 2013; Castillo *et al*. 2024). Consequently, LCs play a vital role in the development of the male reproductive tract, the maintenance of proper spermatogenesis, and overall reproductive function. While LCs dysfunction can lead to various testicular pathologies, their functionality is also notably vulnerable to aging and exposure to exogenous chemical compounds that can trigger cellular death (Svechnikov *et al*. 2010; Wang *et al*. 2017).

Cell death is an essential regulatory feature in multicellular organisms, and mitochondria are central to executing and governing this process (Yin *et al*. 2013; Bock & Tait 2020). Mitochondrial metabolism is crucial for normal energy and biomolecule production, and its dysregulation is a hallmark of numerous pathologies. Crucially, cellular metabolism and death pathways are deeply intertwined in the phenomenon of ferroptosis, an iron-dependent form of regulated cell death driven by excessive lipid peroxidation (Dixon *et al*. 2012; Stockwell *et al*. 2017; Jiang *et al*. 2021). Discovered over the last decade, ferroptosis has emerged as a critical mechanism in multiple diseases and functions (Jiang *et al*. 2021). Considering the central role of mitochondria in regulating cell survival and the intimate relationship between ferroptosis and cellular metabolism, it is postulated that mitochondria play a key role in ferroptotic cell death (Gao *et al*. 2019).

A variety of pharmacological agents, ranging from experimental compounds like Erastin, RSL3, and buthionine sulfoximine to clinically approved drugs such as sulfasalazine, sorafenib, and artesunate, have been identified as potent triggers of ferroptosis. These effects have been extensively documented in both neoplastic models and healthy tissues, including fibroblasts, T cells, neurons, and renal tubular cells (Xie *et al*. 2016). At the molecular level, cysteine availability, glutathione (GSH) biosynthesis, and the proper function of glutathione peroxidase 4 (GPX4) are critical for maintaining ferroptosis in low basal levels (Seok *et al*. 2014; Koppula *et al*. 2020). GPX4, acts as a monomeric glutathione peroxidase that uniquely reduces hydroperoxide groups within membrane-bound phospholipids. By halting lipid peroxidation through this mechanism, the enzyme fulfills a critical biological function (Cozza *et al*. 2017). More recently, lipid metabolism enzymes have been recognized as key pro-ferroptotic drivers. Specifically, factors involved in polyunsaturated fatty acid (PUFA) metabolism dictate cellular lipid composition, embedding peroxidation-susceptible PUFAs into cell membranes (Kagan *et al*. 2017). In this context, the enzyme acyl-CoA synthetase 4 (ACSL4), localized in mitochondria-associated membranes (MAMs) and peroxisomes, acts as a pivotal executioner of ferroptosis by shaping the lipidome and dictating cellular sensitivity to this death mechanism (Doll *et al*. 2017). Furthermore, pathological iron overload, a necessary condition for ferroptosis, has been linked to the deterioration of testicular function, causing spermatogenic disorders and LCs impairment (Yuan et al., 2023).

While iron is essential for normal testicular function and steroidogenesis, iron excess in testicular tissue is a known catalyst for oxidative stress and reproductive dysfunction (Liu *et al*. 2022). Despite the recognized threat of iron-induced damage, the precise dynamics of ferroptosis within LCs and how it intercepts the primary function of these cells, steroidogenesis remains largely unexplored. Therefore, in the present study, we aim to investigate the induction of ferroptosis in LCs and its direct relationship with their steroidogenic capacity. Using both the MA-10 murine Leydig cell line and rat primary interstitial cell cultures, we evaluated the cellular response to ferroptosis inducers (Erastin and RSL3) and explored the transcriptional modulation of *Star*, *Acsl4* and genes related to ferroptosis and the redox cellular state. Furthermore, we investigated the potential protective role of hormonal stimulation with human chorionic gonadotropin (hCG) against ferroptotic damage, providing novel insights into the interplay between cell death mechanisms and endocrine regulation in the male reproductive system.

## 2. Materials and Methods

### 2.1 Reagents and Chemicals

DMEM/F12, gentamicin and 8-bromoadenosine 3′∶5′-cyclic monophosphate (8Br-cAMP) were purchased from Sigma–Aldrich (St. Louis, MO, USA). Penicillin-streptomycin, trypsin solution ethylene diamine tetra-acetic acid (EDTA), and Dulbecco’s modified Eagle’s medium (DMEM) were provided by Gibco-Life Technologies Inc. (Gaithersburg, MD, USA). Fetal bovine serum (FBS) was from PAA laboratories GmbH (Pasching, Austria). Purified hCG was kindly provided by Dr. Parlow (National Hormone and Pituitary Program, National Institute of Diabetes & Digestive & Kidney Diseases (NIDDK, NIH, Bethesda, MD, USA). Horse serum heat inactivates was purchased from Sigma Aldrich-Merck (Darmstadt, Germany). PCR primers were purchased from Thermo Fisher Scientific (Waltham, MA, USA), Macrogen Inc. (Seoul, South Korea) and Integrated DNA Technologies (Coralville, IA, USA). The ferroptosis inducer Erastin was purchased from Cayman Chemical Company (Ann Arbon, MI, USA) and RSL3 from Sigma Aldrich (St.Louis, MO, USA). MitoTracker Red 580 was obtained from Invitrogen (Molecular Probes, Eugene, OR, USA). The 3-(4,5- dimethylthiazol-2-yl)-2,5-diphenyltetrazolium bromide (MTT) reagent and BODIPY™ 581/591 C11 lipid peroxidation sensor were purchased from Invitrogen (Carlsbad, CA, USA).

### 2.2 Cell Culture

The mouse MA-10 Leydig tumor cell line was a gift from Dr. M. Ascoli, College of Medicine (Iowa City, IA) (Ascoli 1981). MA-10 cells were cultured in DMEM/F12 medium supplemented with 20mM HEPES, 15% heat-inactivated horse serum and 50 µg/ml gentamicin. Treatment of MA-10 cells with 8Br-cAMP or hCG was performed in serum-depleted DMEM/F12 for the indicated time points, maintained at 36.6 °C in a humidified atmosphere containing 5% CO_2_.

Primary testicular interstitial cells were isolated from young male Wistar rats (age: 2 months). Briefly, decapsulated testes were subjected to enzymatic digestion using 0.5 mg/ml of type II collagenase. Isolated interstitial cells were plated in low glucose DMEM supplemented with 10% fetal bovine serum (FBS), 1% penicillin/streptomycin, and 1% Amphotericin B and allowed to attach before treatments. All animal procedures were approved by the Institutional Animal Care and Use Committee (CICUAL) of Facultad de Medicina, Universidad de Buenos Aires, resolution (CS) n°: 4081/04.

### 2.3 Cell Viability Assay (MTT)

Cell viability was assessed using the MTT reduction assay. MA-10 cells and primary rat interstitial cells were seeded in 96-well plates at a density of 10^4^ cells/well and allowed to adhere overnight. Cells were then exposed to varying concentrations of Erastin (including 5 µM) and RSL3 (including 10 µM) in the presence or absence of hCG (20 ng/ml) for 24h. DMSO 0.2% (vehicle) was used as control. After treatment, MTT solution (0.5 mg/ml) was added to each well and incubated for 2.5 h at 37°C in the dark. The formazan crystals were dissolved in DMSO. The absorbance at 570 nm was determined using a multi-detection microplate reader, Synergy HT, Biotek (Winooski, Vermont, USA) (Maloberti *et al*. 2010).

### 2.4 Lipid peroxidation measurement

Ferroptosis-associated lipid peroxidation was quantified using the C11-BODIPY 581/591 fluorescent sensor (Martinez *et al*. 2020). Following treatment, cells were incubated with 2 µM of the probe. Lipid peroxidation was determined by monitoring the fluorescence emission shift from the reduced state (measured at ∼590 nm) to the oxidized state (measured at ∼510 nm).

### 2.5 MitoTracker red staining

MitoTracker Red 580 (MTR) dye (Invitrogen) was used to assess mitochondrial activity through fluorescence microscopy (Benzo *et al*. 2024). This compound accumulates in mitochondria with active respiration depending on mitochondrial membrane potential (ΔΨm). MA-10 cells were seeded on 24-well plates on coverslips previously treated with poly-l-lysine for 24 h. Cells were then incubated in serum-free medium with MTR at a final concentration of 150 nM for 45 min at 37 °C in the dark. After incubation, MTR was removed from each well, and wells were washed with PBS. Cells were then fixed with a 4 % paraformaldehyde (PFA) and 5 % sucrose solution for 10 min at room temperature and processed for fluorescence signal detection. Samples were analyzed with a Nikon E200 epifluorescence microscope. Images were obtained at equal exposure time and subsequently analyzed with FIJI software. MTR signal was measured as integrated fluorescence density per area.

### 2.6 RNA Extraction and Quantitative Real-Time PCR (RT-qPCR)

For real-time qPCR total RNA was isolated using Tri Reagent following the manufacturer’s instructions (Molecular Research Center, Cincinnati, OH, USA). Extracted RNA was deoxyribonuclease-treated using RNAse-free DNase RQ1 (Promega, Madison, WI, USA). Reverse transcription was performed using total RNA (1 μg) and M-MLV Reverse Transcriptase (Promega). The expression of *Gpx4* (forward primer: 5′-AGC TAG TCG ATC TGC ATG CC −3′, reverse primer: 5′-CCC TTG GGC TGG ACT TTC AT −3′), *Acsl4* (forward primer: 5′-GTGAACGTATCCCTGGACT - ′, reverse primer: 5′-TGCTTCCCTTCTTGATTTTG −3′), *Tfrc* (forward primer: 5′-CAGATATCAGGGATATGGGTCTAAGTC −3′, reverse primer: 5′-CACGAGCGGAATACAGCCA −3′), *Nrf-2* (forward primer: 5′-CGAGATAAGGCCCGATTGCT −3′, reverse primer: 5′-GGGCATACTGACTGGGTGAG −3′) and *Star* (forward primer: 5′-TTGGGCATACTCACAACCA-3′, reverse primer: 5′-CTTGACATTTGGGTTCCAC −3′) were assessed by real time PCR (Mic PCR instrument, Molecular Biosystems, San Diego, CA, USA) using the SYBR Green Master Mix reagent kit (Applied biosystems, Carlsbad, CA, USA). The reaction conditions were: 5-min in one cycle at 95 °C, followed by 40 cycles at 95 °C for 15 s, 60 °C for 30 s, and 72 °C for 30 s. Gene mRNA expression levels were normalized to *Mus musculus* cyclophilin (forward primer: 5′-TAAACACACAGGACCAGGCC −3′, reverse primer: 5′-CCGTGACCTCCCCAAATACC −3′). Real-time PCR data were analyzed and calculating the 2^− ΔΔCt^ value (comparative Ct method) for each experimental sample (relative gene expression).

### 2.7 Progesterone quantification

MA-10 cells were subjected to the respective treatments for 24 h. Following incubation, the conditioned media was collected and utilized for progesterone (P4) measurement. P4 concentration (ng/ml) was assessed via a competitive ELISA (DiaMetra, Perugia, Italy) following the manufacturer’s guidelines. Absorbance was measured using a multi-detection microplate reader, Biotek Synergy HT. Finally, results were normalized to total protein content per well and present as nanograms of P4 relative to milligrams of total protein.

### 2.8 Bioinformatic analysis of clinical transcriptomic datasets

To evaluate the clinical relevance of the ferroptosis pathway in human male reproductive function, two complementary publicly available human testicular datasets were interrogated. First, baseline single-cell type expression of *Acsl4* and *Gpx4* in human Leydig cells was retrieved from the Human Protein Atlas (proteinatlas.org; single-cell type resource), expressed as normalized counts per million (nCPM). Second, single-cell RNA-sequencing data from the Gene Expression Omnibus (GEO) dataset GSE149512 were re-analyzed to compare Leydig cells between adult donors with normal spermatogenesis and patients with idiopathic non-obstructive azoospermia (NOA) (Zhao *et al*. 2020a).

Count matrices for adult donors with normal spermatogenesis (samples LZ003, LZ013, LZ014, LZ015) and idiopathic NOA (LZ017, LZ018, LZ019) were processed in Scanpy v1.12 (Python toolkit for scalable single-cell transcriptomics analysis). Cells with fewer than 200 detected genes or more than 20% mitochondrial reads were removed; counts were library-size normalized (target sum 1e4) and log1p-transformed. Leydig cells were identified using a steroidogenic marker score (CYP17A1, CYP11A1, INSL3, STAR, HSD3B2) that exceeded parallel germ-cell (DDX4, DAZL, SYCP3) and Sertoli (SOX9, AMH, WT1) scores. To control for germ-cell ambient-RNA contamination, cells with a germ-cell transcript score (including PRM1, PRM2, TNP1, SMCP, DAZL, DDX4, SYCP3) above 0.9 were excluded. Differential expression between conditions was evaluated at the donor level (pseudobulk means; two-tailed Welch’s t-test), retaining donors contributing at least 5 Leydig cells. A p-value < 0.05 was considered statistically significant. For functional interpretation, genes significantly up-regulated in NOA Leydig cells (FDR < 0.05), after exclusion of immediate-early/dissociation-associated, mitochondrial and ribosomal genes, were tested for over-representation in curated gene sets (iron homeostasis, oxidative-stress response, ferroptosis/lipid peroxidation, and LH/cAMP signalling) using one-sided Fisher’s exact tests against the set of expressed genes.

### 2.9 Statistical Analysis

Data are presented as the mean ± standard deviation (SD) of at least three independent experiments. Statistical significance was determined using one-way analysis of variance (ANOVA) followed by Dunnet’s or Tuke’s multiple comparison test using GraphPad Prism software version 9.5.1. A p-value of < 0.05 was considered statistically significant. Differences were denoted as * p < 0.05 and ** p < 0.01, *** p<0.001, **** p<0.0001 vs. control or treatments, as specified in the corresponding figures.

## 3 Results

### 3.1 Leydig cells are sensitive to ferroptosis induction by Erastin and RSL3

To investigate the susceptibility of Leydig cells (LCs) to ferroptotic cell death, we evaluated the effects of Erastin and RSL3 on cell viability using dose-response MTT assays. In the MA-10 mouse Leydig cell line, viability was significantly compromised upon exposure to 5 µM Erastin and 10 µM RSL3. DMSO 0.2% (vehicle) was used as control (Fig. 1A). Moreover, primary cultures of testicular interstitial cells isolated from young Wistar rats were evaluated. Notably, primary interstitial cells exhibited a higher sensitivity to ferroptosis induction, showing a drastic reduction in cell viability at much lower concentrations (1 µM Erastin and 1 µM RSL3) compared to the immortalized cell line (Fig. 1B).

**Figure 1.**
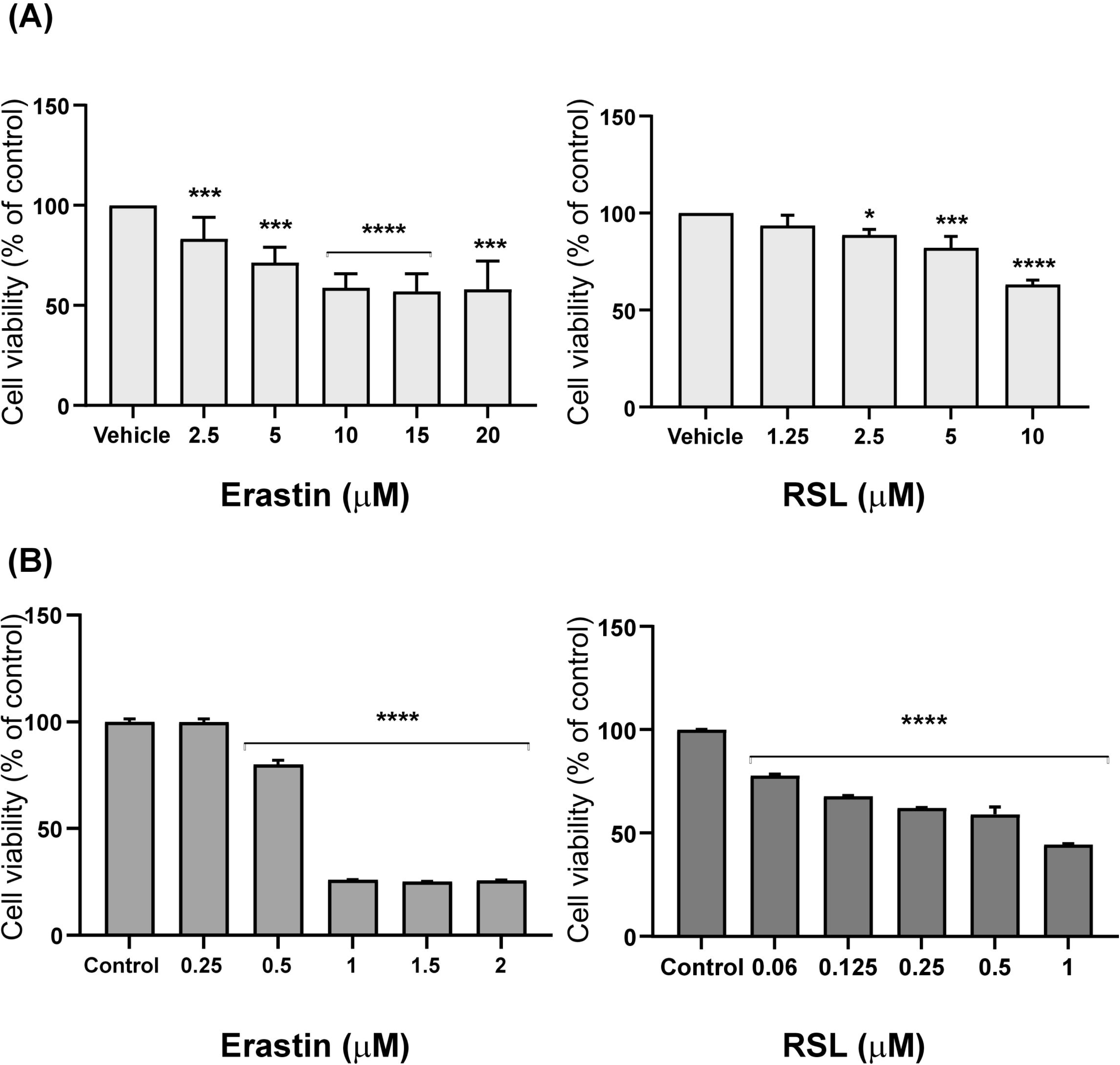
Ferroptosis inducers affect Leydig cells viability. Leydig cells viability was assessed using the ferroptosis inducers Erastin and RSL3 by MTT assay. Dose-response curves were performed. (A) MA-10 cells were treated with increasing concentrations (2.5, 5, 10, 15 and 20 µM) of Erastin (left panels) and (1.25, 2.5, 5 and 10 µM) RSL3 (right panels) for 24 hours. (B) Dose-response curves in interstitial cells with increasing concentration (0.25, 0.5, 1, 1.5 and 2 µM) of Erastin and (0.06, 0.125, 0.25, 0.5 and 1 µM) o RSL3 were performed. Results show a marked sensitivity to both inducers. Data are expressed as mean ± SEM (or SD) of at least three independent experiments. Statistical significance was evaluated by one-way ANOVA followed by Dunnett’s multiple comparison test. (*p < 0.05, ***p < 0.001, ****p < 0.0001 vs. control).

### 3.2 Ferroptosis inducers trigger lipid peroxidation and alter mitochondrial activity

Because the accumulation of peroxidized lipids is one of the defining biochemical signatures of ferroptosis, we quantified lipid ROS accumulation using the BODIPY C-11 sensor. Both Erastin and RSL3 treatments successfully induced lipid peroxidation in our LC models (Fig. 2 A and B). Furthermore, given the central role of mitochondria in LC function, we evaluated mitochondrial activity using MitoTracker Red. Both Erastin (5 µM) and RSL3 (10 µM) treatments significantly increased the MitoTracker Red signal intensity (measured as integrated density per area) compared to DMSO-treated controls (Fig. 2 C and D).

**Figure 2.**
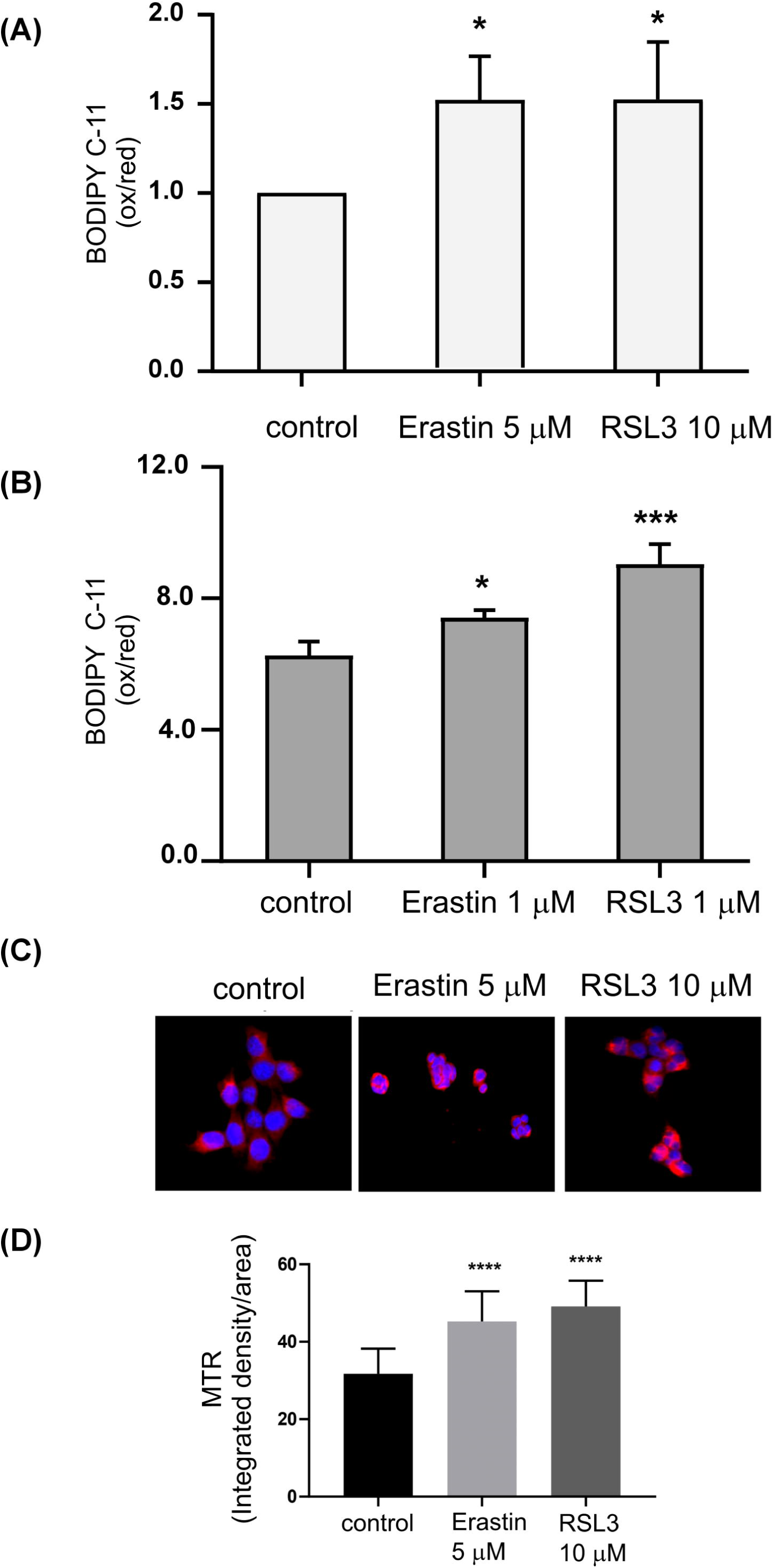
Ferroptosis triggers lipid peroxidation and alters mitochondrial activity. Ferroptosis was induced with Erastin and RSL3 LCs. (A-B) Lipid peroxidation levels in (A) MA-10 cells and (B) interstitial cells following treatment with vehicle (DMSO), Erastin, or RSL3. Analysis was performed using the ratiometric probe BODIPY 581/591 C11, monitoring both reduced and oxidized fractions at their respective excitation/emission wavelengths. Results are presented as fold change of oxidized/reduced relation relative to control (vehicle). Data are shown as the mean ± SD of three independent experiments (C-D) Assessment of mitochondrial membrane potential in MA-10 cells. After treatment with Erastin or RSL3, cells were loaded with 150 nM MitoTracker Red (MTR), a mitochondrial potential-dependent fluorochrome. (C) Representative fluorescence microscopy images showing MTR accumulation (red signal) as an indicator of mitochondrial activity; DAPI (blue) was used as a nuclear counterstain. (D) Quantification of integrated MTR signal as reference of mitochondrial activity. Results are expressed as the mean integrated signal relative to the area analyzed ± SD of at least three independent experiments. Statistical significance was evaluated by one-way ANOVA followed by Dunnett’s multiple comparison test (*p < 0.05, ***p < 0.001, ***p < 0.0001 vs. control).

### 3.3. Erastin and RSL3 differentially modulate ferroptotic gene expression

Given that Erastin and RSL3 promote ferroptosis with different mechanisms, we investigated their effects on several key genes involved in ferroptosis and cellular redox status. We observed that both inducers significantly inhibited *Gpx4* (Fig.3, A) and *Acls4* (Fig.3, B) expression. However, Erastin promotes an increase in *Tfrc* expression in contrast to RSL3, which abrogates the expression of this receptor (Fig.3, C). In addition, our results demonstrate that expression of the antioxidant regulator NFE2L2 (*Nrf2*) is significantly increased following ferroptosis induction by Erastin and RSL3 (Fig.3, D).

**Figure 3.**
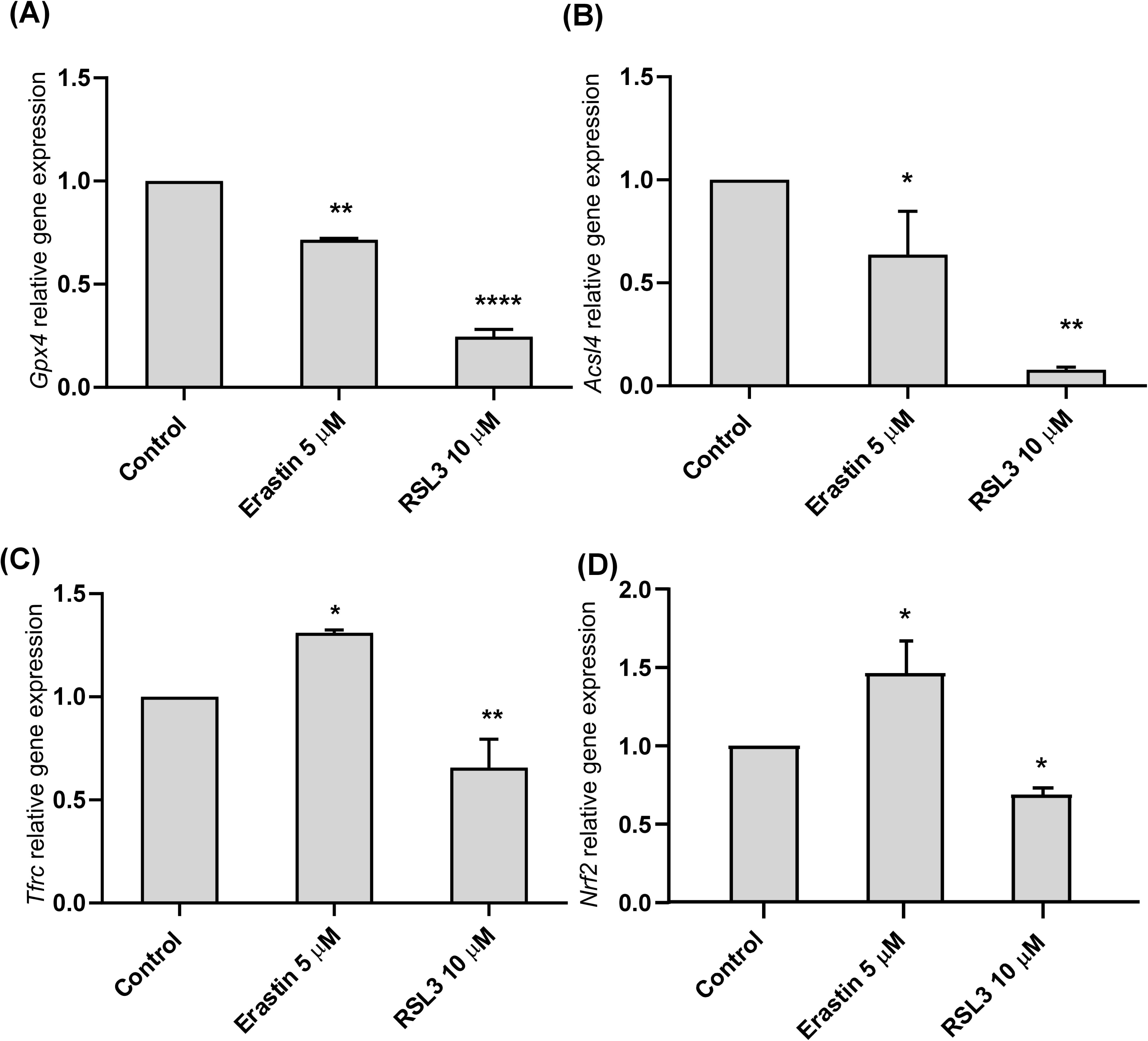
Ferroptosis-related genes are regulated by Erastin and RSL3 in MA-10 Leydig cells. Ferroptosis-related gene expression was assessed through RT-qPCR cells after Erastin and RSL3 treatment in MA-10 cells. (A) Gpx4, (B) Acsl4, (C) Tfrc and (D) Nrf2. Cyclophilin was used as housekeeping gene. mRNA relative expression was calculated as the 2^−ΔΔCt^ value (comparative Ct method) for each experimental sample (relative gene expression). Data represent the mean ± SD of three independent experiments. ANOVA followed by Dunnett’s multiple comparison test. (*p<0.05, **p *<* 0.01, ***p *<* 0.001 vs. control).

### 3.4 hCG stimulation promotes metabolic adaptation and protects steroidogenesis during early oxidative damage

MA-10 Leydig cells viability and function were evaluated under the endocrine control of luteinizing hormone (LH/hCG). The hCG cascade activates the seven transmembrane segments (7TMS) LH receptor and consequently promotes cAMP intracellular increase, PKA activity followed by a plethora of protein phosphorylation processes. We observed that hCG completely mitigates the decrease in cell viability caused by Erastin, maintaining levels comparable to the control. In contrast, hCG fails to protect against RSL3-induced cell death (Fig. 4A). Furthermore, while both ferroptosis inducers significantly decrease basal *Star* expression, co-treatment with hCG significantly rescues *Star* transcript levels in both conditions (Fig. 4B). These results suggest that hormone stimulation participates in the cellular response of LCs to ferroptosis induction. Thus, our findings demonstrate that under the stimulus of hCG, LCs exhibit mechanisms of metabolic adaptation. This hormonal stimulation maintains the functional robustness of the cells, successfully sustaining the induction of the StAR protein during the early stages of oxidative damage.

**Figure 4.**
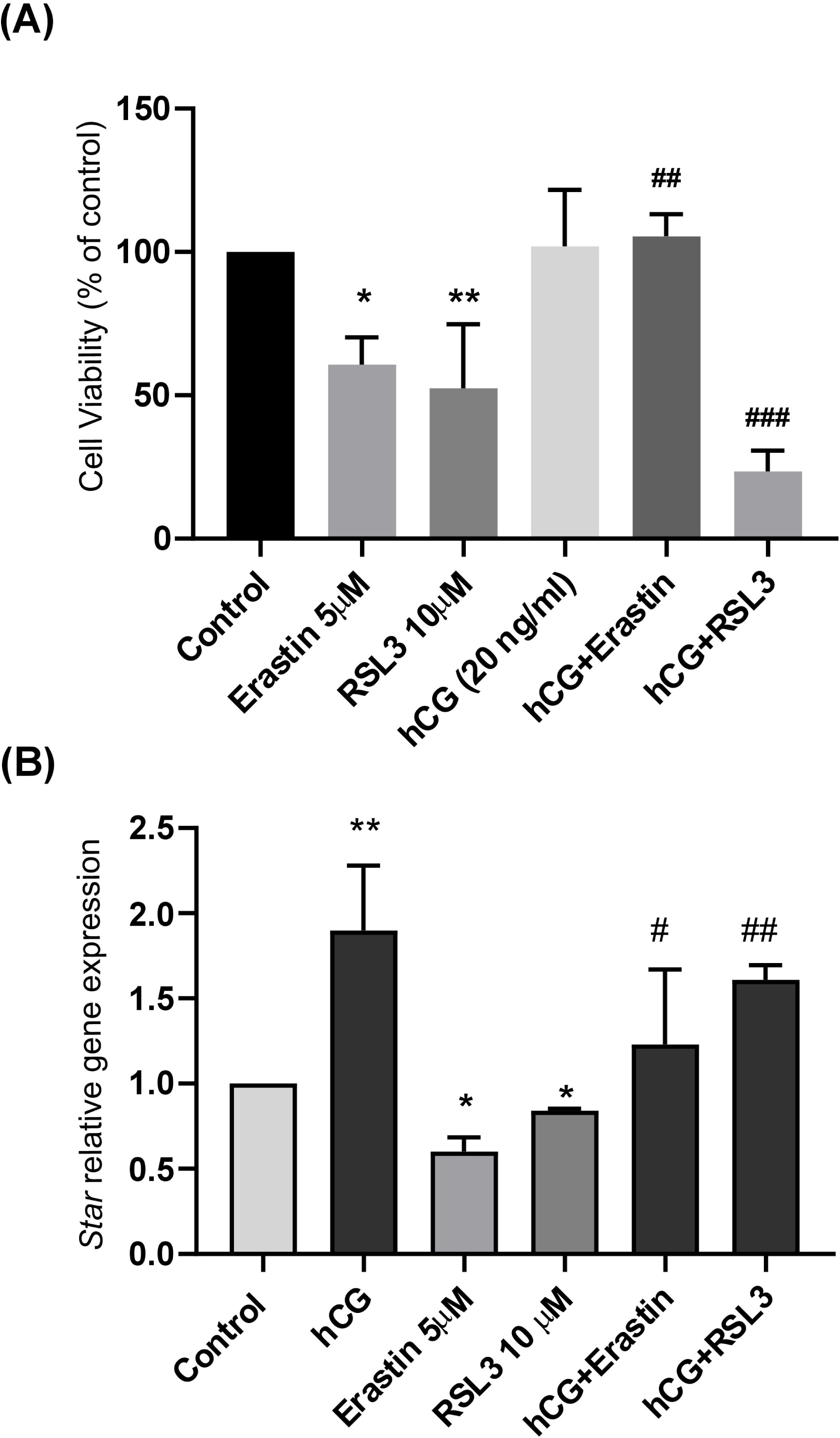
Hormonal stimulation positively modulates cell viability and StAR expression after ferroptosis inducers challenge in MA-10 cells. MA-10 Leydig cells were seeded in 96 well plates. After 48 h, cells were incubated with hCG (20 ng/ml), Erastin (5uM) and RSL3 (10uM) and with or without hCG (20 mg/ml) for 24 h. (A) MTT assay was used to determine cell viability (B) StAR RNA expression levels assessed by RT-qPCR. mRNA relative expression was calculated as the 2^−ΔΔCt^ value (comparative Ct method) for each experimental sample (relative gene expression). Data represent the mean ± SD of three independent experiments. ANOVA followed by Dunnett’s multiple comparison test. (*p < 0.05; **p < 0.01; ***p < 0.001, ****p < 0.0001 vs. control; ^#^ p < 0.05; ^##^p < 0.01; ^###^p < 0.001 vs without hCG).

### 3.5 cAMP-promoted steroidogenesis rescues MA-10 Leydig cells from ferroptosis induction

To further analyze ferroptosis in Leydig cells, we investigated the effect of its induction on both basal and hormone-stimulated steroidogenesis. For this purpose, we used 8Br-cAMP, a widely used cell-permeable analog of the endogenous second messenger cAMP. This compound bypasses ligand-receptor binding to directly mimic the downstream effects of the hCG signaling pathway. The choice of this compound is based on its strong induction of steroidogenesis, as well as its ability to stimulate progesterone (P4) synthesis and secretion into the culture media, particularly in MA-10 Leydig cells. After a 24-hour incubation of MA-10 cells with 5 µM Erastin, 0.25 mM 8Br-cAMP, or a combination of both, P4 concentration in the conditioned medium was evaluated by competitive ELISA as an indicator of steroidogenesis. Although Erastin treatment showed a numerical trend toward decreasing basal P4 levels, a finding consistent with the significant downregulation of StAR expression (Fig. 4B), this difference did not reach statistical significance (Table 1). Treatment with 8Br-cAMP induced a robust, approximately 20-fold increase in P4 secretion. Most importantly, the addition of Erastin did not impair this stimulated response, as co-treatment resulted in P4 levels identical to those of 8Br-cAMP alone (Table 1). This ability to sustain maximal P4 production aligns perfectly with the observation that hCG prevents the downregulation of *Star* by Erastin, confirming that the cAMP pathway effectively preserves the steroidogenic capacity of Leydig cells even during ferroptosis induction (Table 1).

**Table 1.** Effect of Erastin on 8Br-cAMP-stimulated steroidogenesis in MA-10 Leydig cells. MA-10 cells were incubated with 5 µM Erastin, 0.25 mM 8Br-cAMP, a combination of both (Erastine + 8Br-AMPc) and a vehicle (control: 0.2% DMSO) for 24 hours. Progesterone (P4) levels were determined by competitive ELISA and normalized to protein concentration per well. Results are expressed as mean ± SD of at least three independent experiments (ng P4/mg protein). Statistical analysis was performed using one-way ANOVA followed by Tukey’s multiple comparisons test. A significant treatment effect was observed. ANOVA showed a highly significant treatment effect. Different letters indicate statistically significant differences between groups (Tukey’s test, p < 0.05).

| Treatment | Progesterone (ng/mg protein) |
| --- | --- |
| Control | 3.63 ± 0.7<sup>a</sup> |
| Erastin | 0.16 ± 0.03<sup>a</sup> |
| 8Br-cAMP | 71.3 ± 2.7<sup>b</sup> |
| Erastin + 8Br-cAMP | 80.8 ± 27<sup>b</sup> |

### 3.6 Altered expression of key ferroptosis regulators ACSL4 and GPX4 in patients with impaired spermatogenesis

To establish the physiological baseline of the ferroptotic axis in human Leydig cells, we first examined the Human Protein Atlas. In normal testis, Leydig cells constitutively expressed high levels of the ferroptosis-protective enzyme GPX4 (225.3 nCPM) and low levels of the pro-ferroptotic driver ACSL4 (26.4 nCPM), corresponding to an ∼8.5-fold GPX4/ACSL4 ratio favoring protection (Fig. 5A). This configuration is consistent with a ferroptosis-resistant baseline state in healthy Leydig cells. Notably, *GPX4* also showed strong enrichment in spermatids in the same resource, a distribution that confounds whole-tissue comparisons in NOA and justified a cell-type-resolved analytical strategy.

**Figure 5.**
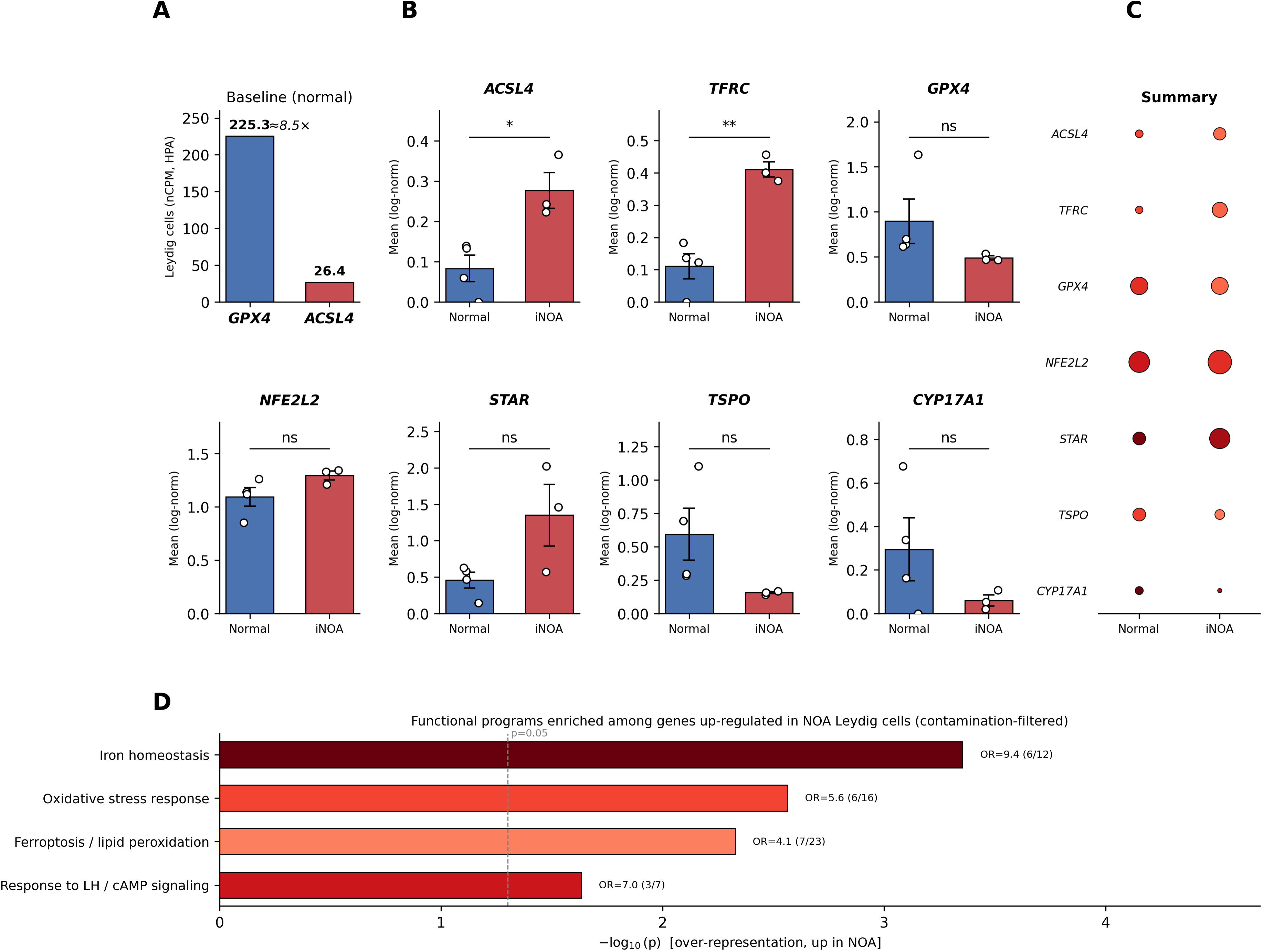
Single-cell analysis of the ferroptotic/iron and steroidogenic axes in human Leydig cells. (A) Baseline expression of GPX4 and ACSL4 in normal human Leydig cells (Human Protein Atlas, single-cell type resource; nCPM); GPX4 predominates over ACSL4 (∼8.5-fold), indicating a ferroptosis-protected baseline state. (B) Single-cell RNA-sequencing analysis of Leydig cells from GSE149512 (4 donors with normal spermatogenesis, 3 NOA donors), after exclusion of germ-cell-contaminated cells. Bars show the mean expression per group (mean ± SEM of donor-level pseudobulk values), with individual donor means overlaid as points; p-values from donor-level Welch’s t-tests. *ACSL4* and *TFRC* are significantly increased in NOA*, GPX4* is concordantly reduced, and the steroidogenic markers *STAR* (up) versus *TSPO* and *CYP17A1* (down) show an uncoupling pattern. (C) Summary dot plot (dot size = percentage of expressing cells; color = mean expression). (D) Over-representation analysis of genes up-regulated in NOA Leydig cells (contamination-filtered) showing significant enrichment for iron homeostasis (OR=9.4, p<0.001), oxidative stress response (OR=5.7, p=0.003) and ferroptosis/lipid peroxidation (OR=4.1, p=0.005); dashed line, p=0.05. *p < 0.05, **p < 0.01; ns, not significant at donor level.

To investigate the clinical and translational relevance of this pathway, we re-analyzed testicular single-cell RNA-sequencing data (GEO dataset GSE149512), comparing Leydig cells from adult donors with normal spermatogenesis and patients with idiopathic non-obstructive azoospermia (NOA) (Zhao *et al*. 2020a). Consistent with previous reports (Tang *et al*. 2022), the relative proportion of Leydig cells was markedly increased in NOA tissue, underscoring why whole-tissue comparisons are confounded by shifts in cellular composition. To additionally control for ambient-RNA contamination from the surrounding germ-cell compartment, cells carrying germ-cell transcripts were excluded, and differential expression was evaluated at the donor level (pseudobulk).

Within Leydig cells, the pro-ferroptotic driver *ACSL4* (p = 0.025) and the iron-import receptor TFRC (p = 0.002) were significantly upregulated in NOA at the donor level, mirroring the pro-ferroptotic shift produced pharmacologically in MA-10 cells (Fig. 5B). *GPX4* showed a concordant reduction in NOA Leydig cells, in the same direction as the *ACSL4* increase and as our *in vitro* findings; this reduction was statistically significant at the single-cell level (p < 0.01) but attenuated to a non-significant trend in the donor-level pseudobulk (p = 0.20), likely reflecting the limited number of normal-spermatogenesis donors available in this dataset. *Nrf2 (NFE2L2)* showed a mild concomitant increase, consistent with an oxidative-stress response accompanying the loss of GPX4-mediated protection.

Notably, steroidogenic components displayed a dissociation pattern in NOA Leydig cells: *STAR* transcript trended upward, whereas *TSPO* and *CYP17A1* trended downward (Fig. 5B, C). This uncoupling, elevated *STAR* transcript alongside reduced downstream steroidogenic machinery, is compatible with sustained gonadotropic drive acting on functionally compromised Leydig cells, in line with the hypergonadotropic hypogonadism characteristic of NOA and with the StAR transcript/function dissociation suggested by our *in vitro* data. Together, these cell-type-resolved clinical data indicate that an imbalance in the ACSL4/GPX4 axis, controlled for cellular composition, accompanies Leydig-cell dysfunction in human male infertility. Consistent with this, over-representation analysis of the genes up-regulated in NOA Leydig cells (after contamination filtering) showed significant enrichment for iron homeostasis (OR = 9.4, p < 0.001), oxidative stress response (OR = 5.7, p = 0.003) and ferroptosis/lipid peroxidation (OR = 4.1, p = 0.005) (Fig. 5D), independently supporting the activation of an iron- and oxidative-stress-linked, ferroptosis-permissive program.

## 4. Discussion

Ferroptosis, an iron-dependent form of regulated cell death driven by excessive lipid peroxidation (Dixon *et al*. 2012), has recently emerged as a pivotal pathological mechanism across a wide spectrum of diseases, including neurodegeneration, ischemic injuries, and cancer (Stockwell *et al*. 2017; Jiang *et al*. 2021). Despite its well-established role in these contexts, its impact on the endocrine system remains poorly understood. Given the high PUFAs content and metabolic demands of testicular tissue, the male reproductive tract is particularly susceptible to this form of oxidative damage (Liu *et al*. 2022). In the present study, we demonstrate that LCs are highly vulnerable to ferroptotic cell death, a process that not only compromises cellular viability but directly disrupts their primary steroidogenic function. Crucially, we unveil a novel endocrine-metabolic crosstalk, revealing that hormonal stimulation via human chorionic gonadotropin (hCG) induces a metabolic adaptation that sustains the steroidogenic machinery during early oxidative stress. These findings provide critical mechanistic insights into the pathophysiology of male infertility associated with testicular iron overload and lipid peroxidation.

We first established the sensitivity of LCs to canonical ferroptosis using Erastin and RSL3, which target distinct nodes of the antioxidant defense system. While the immortalized MA-10 Leydig cell line exhibited a significant, dose-dependent decrease in viability, primary testicular interstitial cultures displayed a markedly higher sensitivity to ferroptotic induction, requiring considerably lower concentrations of the inhibitors to achieve lethal effects. The execution of ferroptosis in our models was confirmed by the rapid accumulation of peroxidized lipids. This heightened vulnerability in primary cultures underscores the physiological relevance of this cell death pathway in the testis. It suggests that endogenous, non-immortalized LCs rely on a highly precise homeostatic balance of iron and lipids, making them exquisitely sensitive to oxidative insults. This intrinsic fragility aligns with clinical observations where uncontrolled lipid peroxidation and iron dysregulation act as primary drivers of testicular dysfunction and male infertility (Liu *et al*. 2022; Guo *et al*. 2026).

Because the rate-limiting step of steroidogenesis, the transport of cholesterol by StAR protein, occurs exclusively at the mitochondria (Stocco 2000; Poderoso *et al*. 2009), we evaluated the impact of ferroptosis on this organelle. Treatment with Erastin and RSL3 resulted in a significant increase in MitoTracker Red signal intensity. In the context of ferroptosis, this functional alteration often reflects early mitochondrial hyperpolarization or swelling preceding cell death (Dixon *et al*. 2012; Gao *et al*. 2019). Our results reveal a profound transcriptional rewiring accompanying mitochondrial distress during ferroptosis, highlighting divergent response mechanisms between Erastin and RSL3. Of particular interest is the marked downregulation of *Acsl4* observed under both treatments, which was most drastic following RSL3 exposure. Given the pivotal role of this enzyme as a pro-ferroptotic mediator, facilitating membrane enrichment with polyunsaturated fatty acids, this strong transcriptional repression may represent a negative cellular feedback mechanism. This could be interpreted as a compensatory survival attempt to limit lipid peroxidation substrates in the face of an imminent lethal insult. The hypothesis of an active cellular rescue effort is further supported by the differential responses observed for *Tfrc* and *Nrf2*. While Erastin induces a significant upregulation of these genes, suggesting the active mobilization of antioxidant defenses and modulation of iron metabolism, the direct inhibition exerted by RSL3 appears to precipitate a more rapid and severe collapse of the core regulatory network, ultimately overwhelming the cell’s capacity to orchestrate a compensatory transcriptional response

Beyond cellular survival, our study highlights the specific vulnerability of the steroidogenic machinery to ferroptosis. We observed that the induction of lipid peroxidation directly impairs the expression of StAR. Interestingly, the mechanistic route of ferroptosis induction dictated the severity of this impairment. Erastin, which triggers GSH depletion via System Xc inhibition (Dixon *et al*. 2012), led to a strong reduction in StAR levels. Conversely, direct Gpx4 inhibition by RSL3 resulted in rapid lipid peroxidation but a comparatively slighter and moderate StAR reduction. This disparity implies that intracellular glutathione levels and broader redox balance might play a more profound role in regulating StAR protein stability or transcription than lipid peroxides alone (Diemer *et al*. 2003), a dynamic that warrants further investigation.

The most striking finding of our study is the protective mechanism elicited by hormonal stimulation. We observed that the presence of hCG actively modifies the cellular response of LCs to ferroptosis. Under physiological endocrine control, hCG stimulation enables LCs to exhibit a remarkable capacity for metabolic adaptation, maintaining functional robustness and successfully sustaining StAR protein induction during the early stages of oxidative damage. This indicates that LH/hCG receptor signaling does not merely drive baseline steroidogenesis, but concurrently activates intracellular defense or compensatory pathways, potentially involving lipid remodeling or enhanced antioxidant capacity, that counteract the accumulation of toxic lipid peroxides (Diemer *et al*. 2003; Maloberti *et al*. 2005; Poderoso *et al*. 2009).

Our viability assays reveal a differential cytoprotective capacity of physiological gonadotropin signaling depending on the node of ferroptosis induction. Strikingly, hCG stimulation effectively rescued LCs from Erastin-induced toxicity yet failed to prevent cell death when ferroptosis was induced by RSL3. Mechanistically, this discrepancy is highly informative. Erastin acts upstream by depleting intracellular glutathione (Dixon *et al*. 2012; Li *et al*. 2020; Zhao *et al*. 2020b), a metabolic deficit that the robust hCG/cAMP signaling axis appears capable of counteracting, likely through the compensatory induction of antioxidant networks. In contrast, RSL3 directly and irreversibly targets the active site of GPX4 (Seok *et al*. 2014). The inability of hCG to bypass direct GPX4 inhibition underscores that the hormone-driven cytoprotective shield is strictly dependent on the functional integrity of this master antioxidant enzyme. Thus, while Leydig cells possess a potent, hormonally regulated defense mechanism against upstream oxidative insults, the ultimate preservation of the steroidogenic machinery relies entirely on basal GPX4 activity to clear ACSL4-dependent lipid peroxides (Doll *et al*. 2017)

Crucially, the physiological relevance of our *in vitro* mechanistic findings is supported by human clinical data at single-cell resolution (Zhao *et al*. 2020a). Re-analyzing testicular single-cell transcriptomes and controlling for germ-cell contamination, we found that the ferroptotic/iron axis is dysregulated specifically within the Leydig-cell compartment of NOA patients, with a significant upregulation of *ACSL4* and *TFRC* at the donor level and a concordant reduction of *GPX4*. In parallel, the steroidogenic machinery showed a dissociation pattern (elevated *STAR* transcript with reduced *TSPO* and *CYP17A1*), compatible with sustained gonadotropic drive acting on functionally compromised Leydig cells. Previous studies have associated ferroptosis with NOA (Hong *et al*. 2024) and documented GPX4-dependent ferroptosis in Sertoli (Bo *et al*. 2025) and germ cells(Yang *et al*. 2025); our analysis extends these observations specifically to the Leydig-cell compartment and, together with our functional data, links the ACSL4/GPX4 axis to steroidogenic function and its hormonal (hCG) protection. We note as a limitation that the number of donors with normal spermatogenesis in available datasets is small, which limits donor-level statistical power (most evident for GPX4); validation in independent single-cell cohorts with larger numbers of normal donors would help confirm these observations. This profound imbalance suggests that the testicular microenvironment in infertile patients is primed for ferroptotic damage, ultimately contributing to the collapse of the steroidogenic machinery.

In the context of Leydig cell biology, our results highlight a fascinating and paradoxical dual role for ACSL4. Physiologically, ACSL4 actively participates in arachidonic acid metabolism to induce StAR protein expression and facilitate mitochondrial cholesterol transport (Maloberti *et al*. 2005; Orlando *et al*. 2013). However, pathologically, ACSL4 is the primary driver of ferroptosis by catalyzing the esterification of long-chain PUFAs into membrane phospholipids, thereby providing the essential lipid substrate for lethal peroxidation (Doll *et al*. 2017). This enzymatic dichotomy suggests that the high basal expression of ACSL4 required for optimal steroidogenesis intrinsically predisposes Leydig cells to a heightened state of ferroptotic vulnerability. When the antioxidant capacity of the cell is compromised, such as upon GPX4 inhibition by RSL3 or during the oxidative stress associated with infertility this ACSL4-driven PUFAs enrichment may transition from a physiological necessity to a pathological liability, potentially triggering mitochondrial membrane collapse and halting steroid production.

In conclusion, our results position ferroptosis as a critical disruptor of male reproductive endocrine function, directly targeting Leydig cell viability and mitochondrial steroidogenic capacity. However, the identification of an hCG-mediated rescue mechanism reveals an intrinsic metabolic plasticity within LCs. Unraveling the exact downstream effectors of this hormonally driven metabolic adaptation will deepen our understanding of testicular biology and could pave the way for novel therapeutic strategies aimed at preserving Leydig cell function and male fertility in the context of oxidative stress and iron-related pathologies.

## Acknowledgments

We thank Silvana Nudler and Marina Vagliente for technical support.

